# Distinct gradients of cortical architecture capture visual representations and behavior across the lifespan

**DOI:** 10.1101/2023.11.29.569190

**Authors:** Xiayu Chen, Xingyu Liu, Patricia Maria Hoyos, Edan Daniel Hertz, Jewelia K. Yao, Zonglei Zhen, Jesse Gomez

## Abstract

The microstructure of cells within human cerebral cortex varies across the cortical ribbon, where changes in cytoarchitecture and myeloarchitecture are thought to endow each region of cortex with its unique function. While fine-scale relative to a cell, these changes at population level impact architectural properties of cortex measurable in vivo by noninvasive MRI, such as the thickness and myelin content of cortex. This raises the question of whether or not we can use these in vivo architectural measures to understand cortical organization, function, and development more broadly. Using human visual cortex as a test bed, we demonstrated two architectural gradients, one in which cytoarchitecture and myeloarchitecture converge and another in which they diverge. These two gradients underlie the structural and functional topography of visual cortex, even predicting the presence of new visual representations. Moreover, the two gradients show distinct visual behavior relevance and lifespan trajectory. These findings provide a more general framework for understanding human cortex, showing that architectural gradients are a measurable fingerprint of functional organization and ontogenetic routines in the human brain.

## Introduction

A fundamental goal of brain science is to elucidate the functional properties of the structural elements of the brain, at an appropriate organizational scale. Classical architectural brain maps including cytoarchitectonic^1,2^ and myeloarchitectonic^3^ maps, derived from postmortem brain sections, have revealed strong correspondence with the functional properties of the cerebral cortex^4,5^. Recent observations of spatial gradients in gene expression across human cortex^6,7^, especially in genes controlling the shape and distribution of dendrites and myelin, also suggest that functional differences across cortex are linked to broad changes in architectural properties^8^. However, these architectural maps are often examined in isolation, lacking interrogation across different maps and individuals. It remains uncertain whether any common or distinct patterns exist among these maps, and if they can offer more reliable and sensitive indicators of functional, behavioral, and developmental relevance. Technological advances in MRI have made it possible to map architectural correlates in human cortex in a noninvasive and individual-specific way, and to test whether any key architectural features of the cortex reflect its functional organization^9–11^. Leveraging the field’s deep understanding of functional organization of visual cortex relative to cortical folding^12–19^, we focus here on visual cortex as a test bed to understand how the architectural patterns of the cortical mantle relates to changes in function. More explicitly, to what extent individuals demonstrate shared architectural features of visual cortex which could be derived from multiple kind of architectural measurements; and how might these structural patterns differ in development and capture differences in visual function and behavior^11^?

Answering such a question would require a large-scale, multimodal MRI dataset to appropriately capture the architectural variations at the level of the population. To that end, we combine three datasets from the Lifespan Human Connectome Project (HCP) which together sample the human lifespan from 5 to 100 years of age^20–22^, and ask if there are shared motifs in architectural features of cortex across individuals and development. Based on T1-weighted (T1w) and T2-weighted (T2w) images, we produce for each individual two distinct maps: a map of cortical thickness^23^ and a map of the T1w/T2w signal ratio^24^. While the thickness map is thought to be attributable to the organization of neuronal, glial, and neuropil tissue^25–28^, the ratio map is thought to be sensitive to the density of myelinated and neurite structures^24,29^, which we refer to more broadly as tissue density. In the case of visual cortex, general trends along the cardinal axes have been observed in architectural features such as myelination and cortical thickness in adults and infants^30–32^, as well as functional properties of neurons such as receptive field size^33,34^ and temporal sensitivity^35,36^. A model explicitly linking these architectural and functional variations across the cerebrum, one that can generalize to yet-mapped regions of cortex as well as explain behavior and dynamics across the lifespan, would be a steppingstone towards bridging structural and functional properties of the living human brain.

## Results

To extract the latent spatial patterns underlying cortical architecture across both properties and individuals, the thickness and tissue density maps from each hemisphere are concatenated across individuals to perform a spatial principal component analysis (PCA)^37,38^ in which participants are features and cortical vertices are samples. As a result, the concatenated maps were linearly decomposed into a collection of orthogonal principal components, consisting of spatial maps (i.e., scores) and individual weights (i.e., loadings) in pairs. The former explains how the architectural properties change across the cortical sheet on each component and the latter describes how individuals’ maps contribute to each component (Figure 1a). The resulting first two principal components (PCs) describe over 50% of architectural variance across the cortical sheet (Figure 1b). Because they are very similar across the two hemispheres (Figure S1), only data from the right hemisphere are presented here for clarity. The contributions of thickness and tissue density indicate that each PC relied on an integration of the two architectural measures at a given spatial location (Figure 1c), rather than a single feature, thus revealing architectural patterns of visual cortex (i.e., score maps) not visible through a single measure alone (Figure S2).

**Figure 1:**
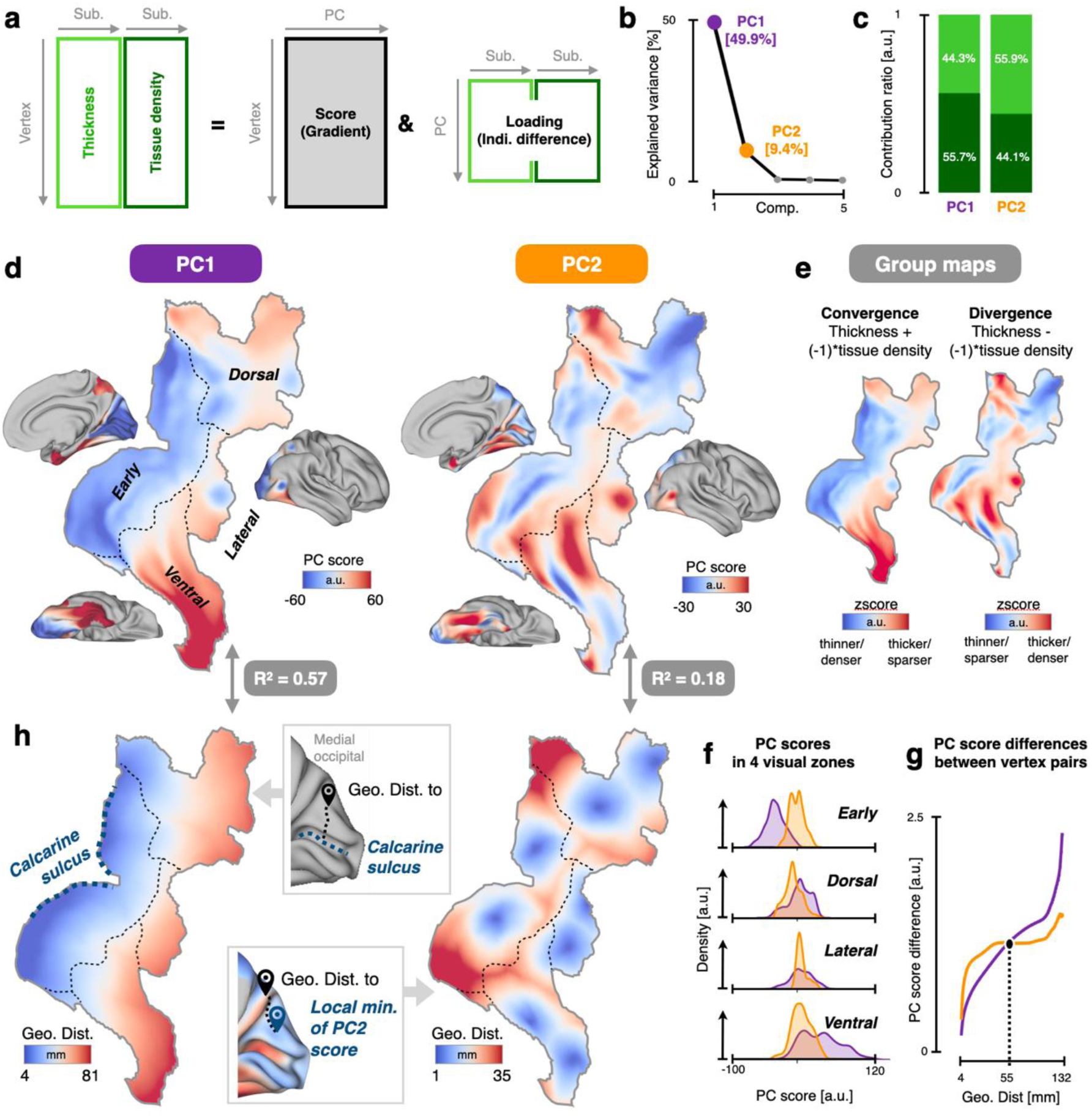
Two architectural gradients scaffold human visual cortex. **(a)** Principal component analysis on the concatenated cortical thickness and myelin/neurite density maps from all participants in HCP-YA to extract architectural gradients of human visual cortex produced a collection of orthogonal principal components, consisting of spatial maps (i.e., score) and individual weights (i.e., loading) in pairs. **(b)** The explained variance ratio of the top 5 principal components (PCs). The first two PCs (i.e., gradients 1 and 2) dominate the explainable variance. **(c)** Contributions of the two architectural measures (thickness and density) to the two gradients. **(d)** Topographic patterns of the two gradients on a flattened cortical surface. Gradient 1 (PC1) displays roughly monotonic change from negative to positive scores across visual cortex, emanating from primary visual cortex V1, while gradient 2 (PC2) showed repeated representation in four localities, mirroring the four visual streams (early, dorsal, lateral, ventral). Black dotted lines: borders where the different visual streams meet, defined using HCP-MMP label boundaries. A.U. is arbitrary units. (**e**) The convergence and divergence between the group average map of thickness and tissue density. **(f)** Histogram depicting gradient scores in the four visual stream regions. Gradient 1 is a large-scale gradient increasing from early to ventral, for example. Gradient 2 is a fine-scale gradient sampled evenly within an individual visual stream. **(g)** The dependence of gradient value differences on geodesic distance are different for the two gradients. Gradient 1 shows larger changes across vertices separated by a long distance, whereas gradient 2 shows larger changes for short distances. **(h)** Geometric models of the two architectural gradients, which were constructed using the geodesic distance of each vertex of visual cortex to specific anatomical landmarks as anchors. The calcarine sulcus and eight local minima of gradient 2 were used as anchors to model gradient 1 and 2, respectively.

Thickness and density maps showed a robust anti-correlation both at the coarse across-area level based on an independent parcellation^39^ and at the finer within-area level, except in primary regions (Figure S3a, b). Given the notable anti-correlation, along with the strong inter-individual correlations within these two sets of architectural maps (Figure S3c), it is anticipated that the first two PCs (Figure 1d) would capture the convergent and divergent patterns between group average maps of cortical thickness and tissue density (Figure S3d). We confirmed this by performing a correlation analysis between PC1 and convergent pattern (thickness map + inversed tissue density map), and between PC2 and divergent pattern (thickness map - inversed tissue density map). As illustrated in Figure 1e, the converging/diverging maps of the group average thickness and inversed density maps are highly correlated with the PC1/2 maps across the visual cortex (both rs = 0.997, ps <0.001). Within-area analyses further confirmed that PC1/2 represent the consistent/deviating components of the thickness–density anti-correlation at finer scales (Figure S3e). This indicates that the PCs have a straightforward and intuitive interpretation. Specifically, PC1 effectively captures the convergence of cortical thickness and tissue density (i.e., how they vary together), while PC2 represents the spatial divergence from this commonality.

Looking into the spatial topography of the two PCs, both PC1 and PC2 maps form spatial gradients whose values change smoothly as one traverses the cortical surface (Figure 1d). Specifically, gradient 1 (i.e., PC1 map) shows an increase in value as one travels from the fundus of the calcarine sulcus either dorsally towards the intraparietal sulcus or ventrally into the anterior temporal lobe. Higher PC1 vertex values correspond to lower tissue density and a thicker cortical sheet (Figure S3d). Interestingly, gradient 2 demonstrates an alternating pattern that fluctuates across cortex. Positive (negative) PC2 scores correspond to higher (lower) thickness or tissue density than expected from PC1 (Figure S3d). Gradient 2 seems to be broadly organized into four distinct zones, mirroring the visual cortex’s division from early visual field maps^40^ into the ventral, lateral, and dorsal processing streams of the visual cortex^41–44^. Quantitatively, the distribution of values for gradient 1 shift across the four processing streams while the value distributions of gradient 2 are evenly sampled within each processing stream (Figure 1f). Furthermore, gradient 2 exhibits a higher spatial frequency in the distribution of its values across cortex. That is, for a given pair of vertices separated by a short distance, gradient 2 tends to show a larger difference in values compared to the more spatially homogenous gradient 1 (Figure 1g). It should be noted that maps of cortical thickness had any variance explainable by curvature removed to account for known thickness differences between gyri and sulci, ruling out the possibility that the alternating architectural pattern was caused by the folding of the underlying cortical sheet. Collectively, these findings demonstrate that gradient 1 forms a large-scale, low-spatial-frequency gradient spanning the entire visual cortex, whereas gradient 2 exhibits a fine-scale, high-spatial-frequency gradient across subregions of the visual cortex.

To get a deeper understanding of the shape of these topographies, we produced simulated models using cortical geometry for the two spatial gradients. For gradient 1, the calcarine sulcus was used as the fiducial line, and vertices of the cortical surface were assigned values based on their minimal geodesic distance to the calcarine sulcus. This simple simulation was able to capture 57.1% of the explainable variance in the topography of gradient 1 (Figure 1h, left), similar to prior distance-based models of this well-known gradient^6^. Gradient 2, which was more complicated in shape, could nonetheless be simulated using anchor points positioned at local minima within each visual processing stream (Figure 1h, right), and vertices of the cortical surface were assigned values based on their minimal geodesic distance to these anchor points. This map of geometric distance also captured a sizeable portion of the variance (17.7%) within the gradient 2 map. A split-half cross-validation yielded similar results, with anchor points and geodesic distances derived from one half of the sample successfully predicting PC2 in the other half (Figure S4), indicating reliable geometric features underlying the spatial organization of PC2. These simulation results again highlight the large-scale and fine-scale characteristics of the two gradients.

Given that gradient 1 values show smooth changes as one travels from early to high-level visual cortex, we hypothesize that it will recapitulate the hierarchical organization of functional features across visual cortex. To test this, we examined the spatial similarity between the architectural gradients and the population receptive field (pRF) properties as measured by the HCP 7T retinotopy dataset^45^. The pRF represents the portion of visual space in which a stimulus evokes a response in a given voxel, and pRF size increases along the visual processing hierarchy^46–48^. We found that gradient 1 was highly correlated with pRF size (Figure 2a) and the known ranking of visual field maps across cortex (Figure 2b), while gradient 2 was not. Strikingly, gradient 1 explains functional features of visual cortex (receptive field size, hierarchical rank) better than the gradient derived from resting-state functional connectivity (RSFC) within visual cortex (Figure S5a-c). Further evidence for this hypothesis can be found when examining temporal properties of visual cortex function. It is widely recognized that the frequency at which the BOLD signal fluctuates during resting-state functional MRI^36,49,50^ is different in distinct regions of cortex. This temporal property of the BOLD signal, quantified as a fractional amplitude of low-frequency fluctuation (fALFF)^51^, correlates well with temporal properties of receptive fields in visual cortex^35,36,52^. Here, we find that this temporal gradient as measured by fALFF is well-described by gradient 1 in a simple regression analysis (Figure 2c).

**Figure 2.**
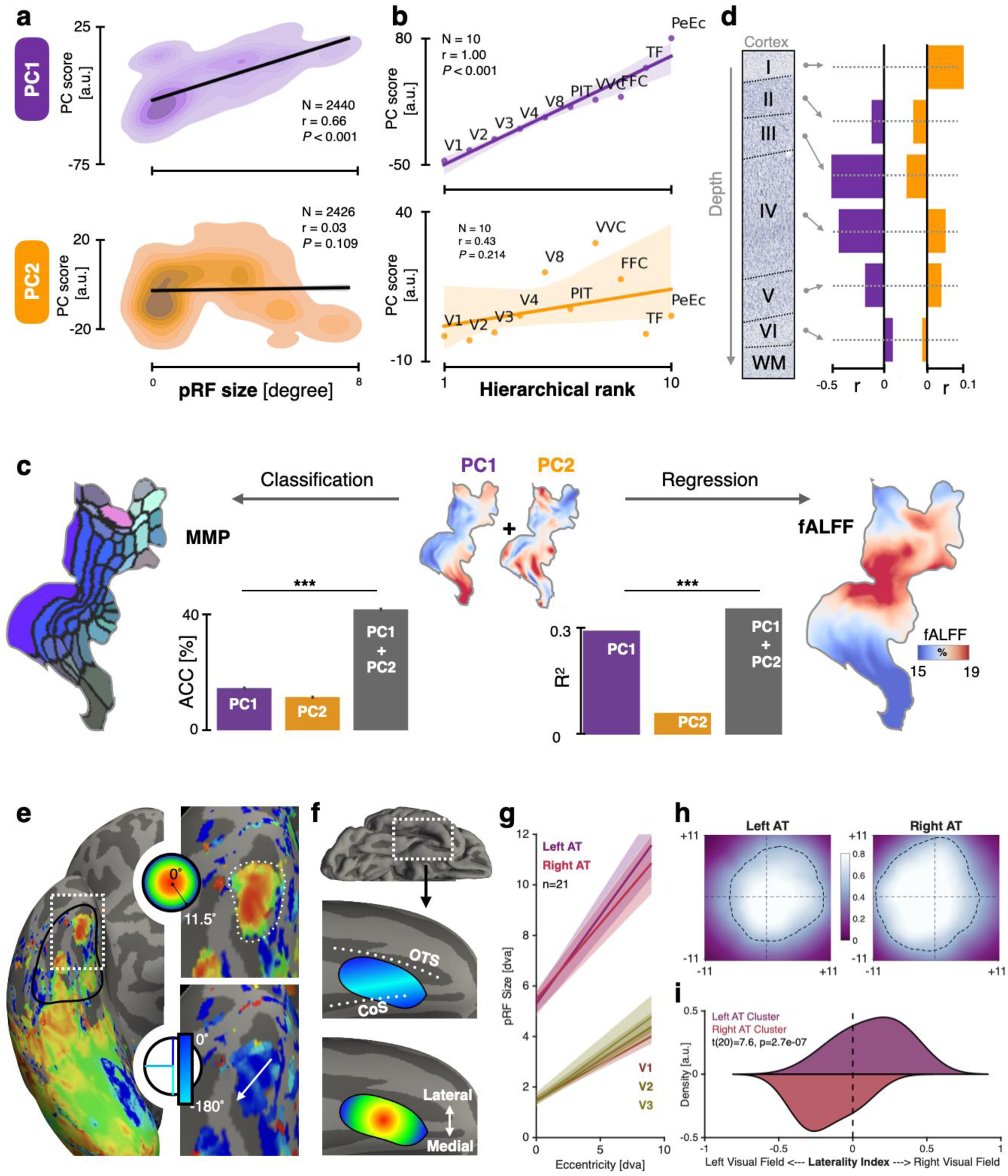
The functional and microstructural properties of the two architectural gradients. **(a)** Gradient 1 was highly correlated with the pRF size (r = 0.66), while the gradient 2 was not (r = 0.03). **(b)** Gradient 1 was perfectly correlated with the hierarchical rank of the 10 visual areas within the ventral visual stream (Spearman rank ρ = 1.00), while gradient 2 was not. **(c)** The functional significance of the architectural gradients was evaluated by measuring to what extent each gradient is related to areal differentiation of the visual cortex (left) and the global functional organization measured by fractional amplitude of low-frequency fluctuation (fALFF) from resting-state fMRI (right). The combination of gradients 1 and 2 greatly improved the predictive power of classifying visual areas compared to using either gradient alone. However, gradient 1 contributes more than gradient 2 in predicting the large-scale functional organization (i.e., fALFF map). **(d)** Cell body density from the BigBrain dataset is quantified for each cortical layer at each vertex and correlated with each gradient map. Gradient 1 was mainly correlated with cell body density in Layers III and IV, while gradient 2 was mainly correlated with cell body density in Layer I. **(e)** The architectural gradient 2 predicts the presence of novel visual representations in the anterior temporal lobe. Left: example participant with the pRF eccentricity map displayed on the inflated cortical surface. The highlighted region (white dotted line) is the subject of the zoomed insets on the right. The black outline delineates the anatomical region from which pRF data was extracted. The putative visual representationcluster is outlined on the insets, showing a radial eccentricity representation, and a perpendicular representation of polar angle travelling roughly medio-anterior to latero-posterior as indicated by the white arrow. **(f)** Illustration on an inflated cortical surface illustrating that the AT-cluster of retinotopic maps is located near the anterior intersection of the occipitotemporal (OTS) and collateral sulci (CoS). The AT-cluster representations demonstrate perpendicular representations of pRF eccentricity and polar angle. **(g)** In all 21 participants, pRF size and eccentricity from all above-threshold vertices within the anatomically defined region (black solid line from panel e) are extracted, binned by eccentricity, averaged across participants, and then lines-of-best fit are modeled across the averaged data. Shaded regions represent bootstrapped 68% confidence intervals. (**h**) Average visual field coverage maps of left and right AT voxels. Lighter colors represent higher coverage density, black outline denotes 50% pRF density compared to maximum. Value of 1 denotes all participants had maximum pRF coverage at that point in space. (**i**) Kernel density histogram denoting laterality index across participants. Value adjusted such that +1 denotes purely right visual field coverage across an individual’s pRF fits, -1 denotes purely left laterality, and 0.5 is bilateral coverage.

Thus, the convergence relationship between cortical thickness and density across visual cortex appears related to broad changes in functional features. In contrast, gradient 2 appears to act more locally, showing greater spatial inhomogeneity with interdigitating peaks and valleys of scores within individual visual streams. We hypothesized that these local deviations from the canonical thickness and density of cortex underlie the finer-scale division of visual cortex into categorically distinct regions. That is, does the arealization of cortex into distinct regions involve these regions becoming more distinct from a typical cortical sheet (i.e., gradient 1)? To test this, we hypothesize that gradient 2, when combined with gradient 1, will improve the differentiation of cortical regions as defined by the HCP multimodal parcellation^8^. Through a classifier-based analysis, regional classification was significantly higher when gradient 2 is combined with gradient 1 (Figure 2c). Additionally, the two architectural gradients outperform the first two gradients derived from resting-state functional MRI in a similar classification approach (Figure S5d).

Lastly, if the two architectural gradients truly relate to the functional organization of visual cortex in distinct ways, then these differences should be mirrored in the way cytoarchitecture contributes to each structural gradient. Based on the BigBrain dataset^53^, we extracted cell body density data for each of the six cortical layers to produce six structural maps projected onto the cortical surface. For gradient 1, we find that changes in its structural features correlate strongly with cell density of layers III and IV, where the pronunciation of layer IV decreases with increasing distance from the calcarine sulcus. Gradient 2 was most correlated with cell density of layer I, with positive scores overlapping cortex with thicker superficial layers of cortex (Figure 2d). Because layers III and IV are primarily involved in feedforward connections whereas layer I plays a predominant role in feedback projections, this finding might suggest that gradient 2 is a structural fingerprint of differential feedback connectivity in visual cortex.

Given the relatively tight correspondence between these architectural gradients and the functional organization of visual cortex, can this structural-functional coupling generalize to regions of visual cortex that have not yet been mapped? Upon examination of the topology of gradient 2, a pattern emerges between the gradient and retinotopic representations. While most of the anchor points for the gradient 2 map simulation correspond to visual field map clusters which share a foveal representation (V1-V4, VO1-2, IPS0-1, IPS2-3, TO1-2)^40,54^, one additional anchor appeared in the anterior temporal lobe near the location where the occipitotemporal sulcus (OTS) merges with the collateral sulcus (CoS) more medially (Figure 1d,h). If we assume a correspondence between gradient 2 anchors and visual field map clusters, then this anterior-most anchor would suggest an additional cluster of visual field maps in the anterior temporal lobe, one which has not yet been described in the literature. To test this hypothesis, and potentially demonstrate the predictive power of these architectural gradients to unmapped cortex, we performed pRF mapping^46^ on 21 participants with high-contrast, socio-ecological images to better drive neurons of high-level visual cortex often tuned for such complex objects^55^.

We indeed find a cluster of visually responsive pRFs in the anterior temporal lobe located medially, overlapping the CoS and extending laterally towards the OTS usually just beyond the anterior tip of the fusiform gyrus but sometimes overlapping it (Figure 2e). This location is consistent with a previous report of face-selectivity in the anterior temporal lobe^55^ and is located within perirhinal cortex and approximately located at the border between Brodmann areas 35 and 36^56,57^. These maps were observable in the majority of hemispheres (see Figure S6). This cluster of maps shows a clear radial representation of pRF eccentricity, with voxels near the center of the cluster sampling central visual space, and those near the outer boundary of the cluster sampling peripheral visual space. Perpendicular to this radial eccentricity representation was a representation of polar angle, with two upper visual field representations separated by a shared lower visual field representation, usually oriented at an oblique angle to the CoS but sometimes parallel with it (Figure 2e, f). A hallmark feature of visual pRFs is that they increase in size as one ascends the visual processing hierarchy, and the positive relationship between pRF eccentricity and size tends to become more dramatic as well^33,46–48^. To test this, we extract pRF fits from vertices with variance explained greater than 10%. We find that consistent with its high position within the processing hierarchy, pRF sizes are significantly larger than in earlier visual field maps V1 through V3, and the linear function relating pRF eccentricity and size yielded larger slopes compared to V1-V3 (Figure 2g). Consistent with spatial computations in earlier visual field maps, these anterior temporal (AT) representations, display pRF coverage that mainly samples the contralateral visual field^15,34,47^, although it was not uncommon for pRF centers to sample ipsilateral visual space (Figure 2h). To quantify if the visual field sampling in each AT representation was significantly contralateral, we derived a laterality index^58^ for each individual left or right AT cluster and display the histogram of indices (Figure 2i). We find a significant difference in laterality indices between left and right AT pRFs (paired-samples t-test, t(20) = 7.6, p = 2.7 × 10^-7^).

If these architectural gradients are capable of extrapolating to functional representations in visual cortex, to what extent can they also describe the behaviors supported by visual cortex? In addition to providing spatial maps of scores, the PCA provides a weight or sense of fit describing how a given participant relates to a given gradient. To answer the question above, a canonical correlation analysis (CCA)^59^ was performed to examine how individual participant weights for the two gradients can predict the individual behavioral performance from 15 vision-related behavioral tasks^60^ (Figure 3a). As shown in Figure 3b, both gradients show significant correlation with visual ability. However, in comparison to the architectural convergence pattern (gradient 1), the divergence pattern (gradient 2) demonstrates a stronger correlation with visual behaviors. Moreover, the two gradients are associated with distinct and unrelated behavioral profiles (Figure 3c): Behavioral variables related to attention, visual acuity and inhibitory control are more closely associated with gradient 1, while vocabulary comprehension, fluid intelligence, spatial orientation processing ability, and nonverbal episodic memory ability are more strongly linked to the gradient 2. The distinct behavioral mapping profiles of the convergence and divergence gradients further underscore each architectural gradient’s unique contribution to brain function and highlighting the cognitive importance of the architectural gradients.

**Figure 3.**
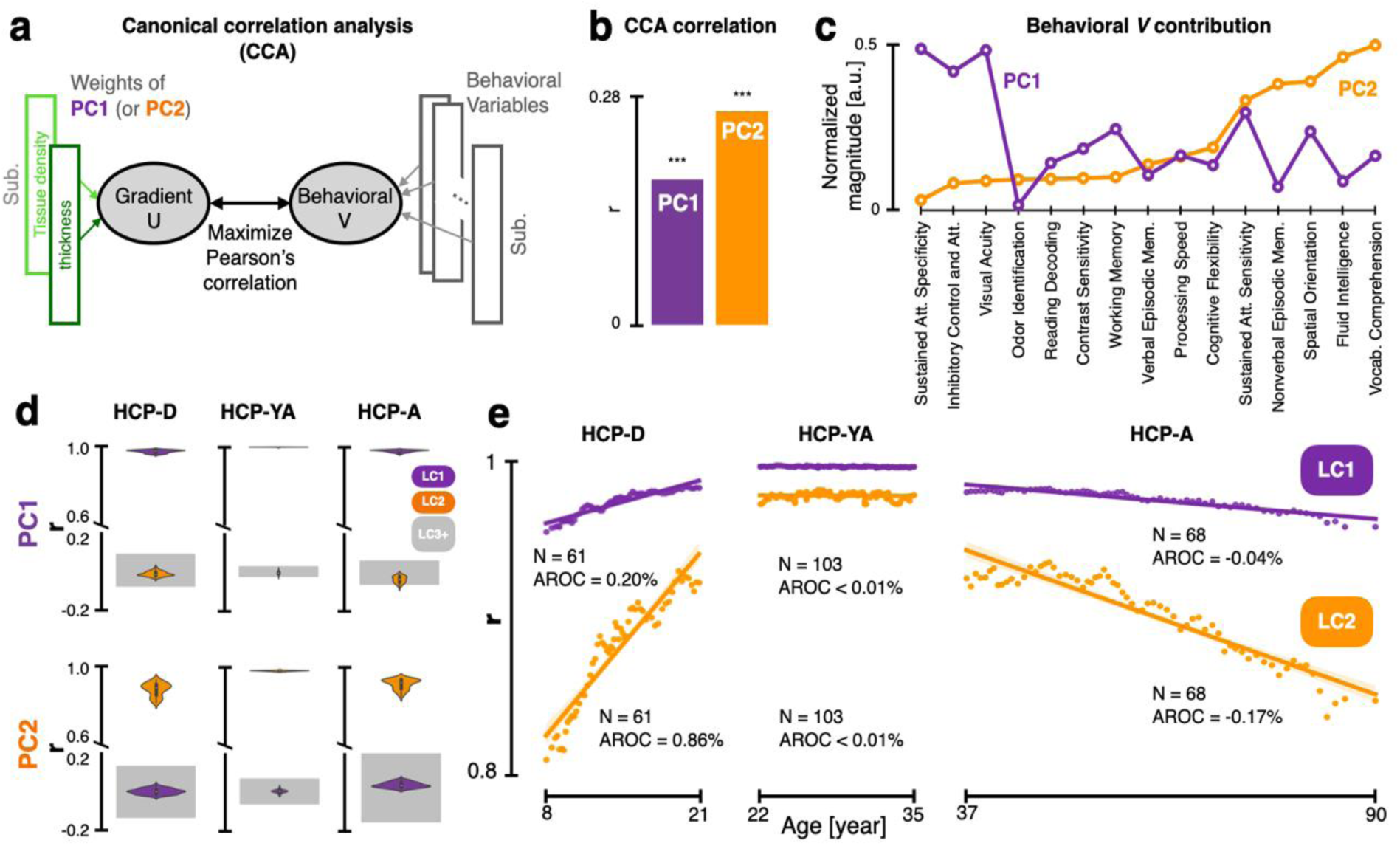
The relevance of the architectural gradients to visual behavior, development, and degeneration across the lifespan. **(a)** Canonical correlation analysis (CCA) was used to associate multiple vision-related tasks with the two weight-vectors of each architectural gradient in the HCP-YA. CCA finds the linear combination of variables that best associate measures from the two data domains across participants. **(b)** Both architectural gradients can significantly predict individual visual ability. However, gradient 2 showed stronger correlation with visual behaviors than gradient 1. **(c)** The normalized magnitude of behavioral factor weights from CCA indicate that gradient 1 was correlated more with low-level visual abilities, while gradient 2 was correlated more with higher-level visual abilities. **(d)** Sliding window spatial PCA was performed across the lifespan (PCA on a given age-bin results in “lifespan component”, LC) to compare how the patterns of gradient 1 and 2 change with the age. The top two LCs (i.e., spatial maps) extracted from each age window correlate strongly with their respective component from the HCP-YA, confirming that the architectural gradients can be observed across the lifespan. **(e)** Correlations between LCs and PCs reveal that gradient 2 shows more development and degeneration across childhood and aging, compared to gradient 1.

If these architectural gradients across the cortical sheet correspond to differences in cortical tissue content, brain function, and behavior, as evidenced above, then they should also change across the lifespan, given that behavior and neocortical tissue structure develop dramatically during childhood across visual cortex^27,32,61^. The two spatial gradients described above were derived from the young adult dataset (HCP-YA). We can therefore repeat the spatial component analysis at various stages of the lifespan. We can ask if the first two principal gradients replicate in these separate stages, and determine how they may change, if at all, across the lifespan using HCP development (HCP-D) and aging (HCP-A) datasets^62^. We binned participants in equal-sized windows of increasing age, deriving within each window the top PCs and correlating their score maps with that from the young adult dataset. To distinguish the PCs of each age window from the PCs of the young adult dataset, we referred those derived from developmental age-bins as lifespan components (LC). We found that LC1 and LC2 from the developmental (n = 652 participants, 351 females, ages 5-21), adult (n = 1070, 586 females, ages 22-37), and aging (n = 725 participants, 406 females, ages 36-100) data at every age bin show a high correspondence to gradient 1 and gradient 2 from the young adult data respectively, compared to other LCs (Figure 3d). This demonstrates that the two gradients derived from lifespan data at each window replicate those of the young adult dataset, allowing us to trace their developmental trajectories.

Examining the correlation between the young adult gradient and LC within each developmental window, we first found that LCs, as expected, are stable during young adulthood (Figure 3e, middle). However, we found a linear change across childhood, with the topography of LC1 and LC2 becoming more adult-like with maturation (Figure 3e, left). The trajectory of each LC was unique, with a significant age × LC interaction (F₁,₁₁₈ = 257.01, p < 0.001). LC2 showed a significantly larger developmental effect than LC1, with an annualized rate of change (AROC) four times that of LC1(0.86% vs. 0.20%). Finally, if gradients solidify their structural topography across childhood and adolescence, do they show degeneration in later adulthood? We can make two a priori hypotheses here: first, that both LCs will show linear loss of their adult-like topographies and second, that LC2 will show more rapid degeneration than LC1, consistent with the developmental retrogenesis trends observed in white matter development^63^. As shown in Figure 3e (right panel), the sliding-window gradient analysis on the aging dataset revealed that both hypotheses are supported, LC1 and LC2 both show linear loss of their topographies with LC2 showing more dramatic degeneration than LC1 (LC1 AROC = -0.04%; LC2 AROC = - 0.17%; age × LC interaction F₁,₁₃₂ = 263.85, p < 0.001).

## Discussion

Overall, we provide evidence for two architectural gradients across human visual cortex, each of which captures a distinct relationship between the thickness and tissue density of the cortical sheet. While the first gradient captures a relationship in which thickness and density measures vary in agreement with one another, the second captures a relationship in which these two features diverge from one another. The convergence pattern, arising from the negative correlation between thickness and density, is consistent with previous findings^64–66^ and may support the balloon model, whereby cortical thinning is associated with tangential stretching due to myelination^67^. In terms of cortical spatial organization, the convergence map scaffolds the cortex into a processing hierarchy, while the divergence map between these features appears to contribute to the finer-scale arealization of the cortex into distinct regions. These two architectural gradients together describe well the broader functional landscape and seem to relate to unique aspects of cytoarchitecture across cortex. Where the first gradient correlates strongly with pRF size of the visual system, and is capable of ranking regions into their ground-truth hierarchical ordering, the second gradient appears to differentiate visual cortex into distinct zones within a hierarchical level. The second gradient not only predicted the location of a new contralateral visual representation in anterior temporal cortex, but also demonstrated more dynamic changes across the lifespan which may correlate with visual behavior development.

These data would suggest that gradient 2 tracks the functional differentiation of visual cortex into unique regions, wherein regions occupying the same hierarchical level within gradient 1 show distinct values in gradient 2. The current data offer strong evidence in this regard through two separate observations. First, adding the divergent gradient into a classifier along with the convergent gradient significantly improved classification performance of cortical region labels (Figure 2c). Second, cortical regions where divergent gradient scores transitioned from positive to negative values appeared to be spatially related to the location of shared foveal representations in visual field map clusters. We observed such a transition zone in the structural measures of the divergent gradient in anterior temporal cortex. Consistent with this structure-function coupling, we observed a novel, contralateral visual field representation with a clear radial eccentricity and largely perpendicular polar angle transitions (Figure 2e-i). The organization of polar angle in anterior temporal cortex was not as orderly as earlier visual cortex, but consistent with disorderly polar angle representations in higher-level dorsal visual cortex^68^. Thus, whether these representations constitute formal visual field maps will require higher-resolution pRF mapping, but their observed location provides strong empirical evidence for the predictive power of our cortical model on the functional topography of the brain. These new contralateral visual representations also provide greater context for previous observations of a face-selective region in the anterior temporal lobe^69^, which may neighbor or overlap these novel AT pRF clusters. This relationship would be consistent with previous work showing that face-selective regions on the fusiform gyrus neighbor foveal representations of nearby visual field map clusters in more posterior portions of ventral occipitotemporal cortex^70,71^.

Given that gradient 2 shows the strongest relationship to behavior and tracks developmental changes across the lifespan, it might suggest that the differential development between cortical regions during childhood more strongly drives maturation of visual behavior, compared to broader-scale structural changes. Indeed, previous work has shown that divergent developmental trajectories between category-selective regions of high-level visual cortex underlies behavioral maturation of high-level visual abilities^61^ and that a disruption in this differential structural development may underlie disorders of high-level vision^72^. Future work can examine if functional arealization at earlier developmental timepoints, as in infancy^73^, follows this prediction.

Likewise, our results hold implications for understanding human neurodevelopment. The nearly mirror symmetric development and degeneration of these architectural gradients between childhood and later adulthood offer strong evidence that gradient 2 follows the pattern of retrogenesis^74^ across the lifespan. It is presently unclear if the changes in cortical tissue during childhood that drive the developmental emergence of gradient 2 are the same tissue compartments which degenerate later in life (Figure 3e). Indeed, while our current analyses focus on cortex, the expanse of visual cortex described here is innervated by several major white matter pathways such as the vertical occipital^75^, the inferior longitudinal^76^, and the arcuate fasciculi^77^, all of whose development has been shown to be behaviorally relevant^72,78–81^. Therefore, the extent to which these architectural changes reflect local tissue structure versus connectomic features^82^ should be clarified in future work. Furthermore, while gradient 1 described a typical relationship between the thickness and density of tissue in the cortical sheet, gradient 2 captures how these two features diverge from this canonical relationship and exhibits greater behavioral relevance across a range of visual behaviors. This might suggest that the patterning of cortex into distinct cortical regions necessitates deviation from a canonical cortical sheet, reminiscent of the protocortex hypothesis of gestational neurodevelopment in which largely similar progenitor cells in the primordial cortex differentiate as a result of distinct thalamic inputs across cortex^83–85^. The morphological changes during gestation that lead to arealization of cortex into distinct regions are certainly distinct from the childhood structural measures quantified here through MRI, but may nonetheless suggest that the ontogenetic routines which serve to differentiate cortex into cytoarchitectonically distinct zones during gestation likely have long-term influences on the postnatal development observed here.

It is worth emphasizing that while classical cortical parcellation emphasizes sharp cytoarchitectonic boundaries^1^, the gradients identified here—derived from interindividual variability in microstructural features—do not contradict this view but instead provide complementary information that may capture supra-areal organizational principles describing how distinct cortical regions relate to one another across the cortical sheet.

These findings offer evidence that there are architectural gradients, measurable with MRI, that are shared across individuals. These shared patterns of cortical sheet morphology track the functional organization and computations of the underlying cortical sheet across the human lifespan. These findings provide a normative benchmark for future work examining how deviations from these shared mesoscale architectural patterns underlie neurological disorders.

## Methods

### Human Connectome Project Data

The publicly available data from the HCP Young Adult (HCP-YA)^21,86^, Development (HCP-D) and Aging (HCP-A)^62^ were used in the study. The three large-scale brain imaging studies collect behavioral and multi-modal MRI data in healthy participants from 5 to 100 years of age, and thus provide us with opportunities to characterize brain changes across the human lifespans. Only structural MRI, resting-state fMRI and behavioral data were used in the study. The use and analyses were approved by the Institutional Review Boards of Beijing Normal University (No. ICBIR_A_0111_001) and Princeton University (No. 13074).

After excluding participants with invalid MSM-All registration and those without any resting-state functional MRI (rs-fMRI) data, we obtained multi-modal MRI and behavioral data for 1070 HCP-YA participants (586 females, ages 22-37, S1200 release). For each participant, T1-weighted (T1w) and T2-weighted (T2w) structural images (0.7mm isotropic voxels) and functional images (2mm isotropic voxels; TR=720ms) were acquired on the HCP’s customized 3-Tesla Siemens Skyra scanner using a 32-channel head coil. The rs-fMRI paradigm included two sessions, each session itself including two runs with opposite phase-encoding directions (R/L and L/R, each 15 minutes long). All the structural and functional MRI data were preprocessed using the HCP minimal preprocessing pipelines, and more information regarding data acquisition and preprocessing is available from previous work^86–88^.

The HCP-D and HCP-A datasets were acquired on a 3T Siemens Prisma scanner with similar protocol as the HCP-YA data^20,22^. Structural MRI data (0.8mm isotropic) from 652 HCP-D participants (351 females, ages 5-21) and 725 HCP-A participants (406 females, ages 36-100) were used in this study (Lifespan HCP release 2.0). Preprocessing of these two datasets was nearly identical to that of the HCP-YA with small adaptations to account for the variability of the wider age range^62^. The HCP data used in this study were in fsLR_32k cortical space based on the MSM-All registration^89^, and cortical thickness data used in the study have been regressed out to exclude the linear effect of cortical curvature.

### BigBrain Data

The BigBrain dataset is a volumetric reconstruction (20 μm isotropic) of a histologically processed postmortem brain of a human male 65 years of age. Sections were stained for cell bodies, imaged, and digitally reconstructed into 3D volume^53^. The white and pial surfaces of the BigBrain were extracted at the gray-white matter boundary and gray matter/cerebrospinal fluid (CSF) boundary^90^, respectively. The 3D laminar atlas, including six cortical layers, was also derived at 20 μm isotropic resolution^90,91^. Based on the surface registration to the MRI-based MNI152 template surface, the cytoarchitectural information from each layer of the BigBrain can be linked to in vivo neuroimaging data.

### Population receptive field (pRF) experiment

We performed pRF mapping on 21 participants with high-contrast, ecological images to better drive neurons of high-level visual cortex often tuned for such complex objects. We adapted the experiment used in the HCP 7T Retinotopy Dataset^45^. Stimuli consisted of slowly-moving bar-shaped apertures of 2-degree width filled with a dynamic colorful texture. Textures presented within the bar aperture were updated at a rate of 7 Hz. Textures included randomly-presented cartoon scenes depicting people, animals, characters, text, limbs and objects evenly spanning the width of the stimulus aperture. Participants were asked to fixate on a central dot while attending to the bar, monitoring it for the random appearance of a target cartoon stimulus (a grid of wiggling bumblebees) which appeared for a 500ms duration, 10 times during the experiment. Each run lasted 300s, and participants completed 3 to 4 runs.

### Data Analysis

#### Definition of human visual cortex

Human visual cortex was defined by grouping the 44 visual areas from the HCP multimodal parcellation (MMP) atlas^8^. All of these areas located in the occipital, parietal and temporal cortices and show a significant BOLD response to visual objects. According to the well-established model of the visual cortex, these areas are grouped into four visual processing streams: early stream (V1, V2, V3, V4), lateral stream (V3CD, LO1, LO2, LO3, V4t, FST, MT, MST, PH); dorsal (V3A, V3B, V6, V6A, V7, IPS1, LIPv, VIP, MIP, 7Am, 7PL, 7Pm, IP0, IP1, DVT, ProS, POS1, POS2, PCV), and ventral (V8, VVC, PIT, FFC, VMV1, VMV2, VMV3, PeEc, PHA1, PHA2, PHA3, TF). Please see supplementary information for the detailed descriptions of these areas (Table S1).

#### Extracting architectural gradients of human visual cortex

The thickness of cortex, and the density of neurite and myelin content (referred to here as tissue density), are two widely used mesoscale in vivo architectural measures derived from structural MRI and were used to extract architectural gradients of human visual cortex. Cortical thickness was measured as the shortest distance between each vertex on the white matter surface and the pial surface^23^, while cortical tissue density was measured by the ratio of T1w to T2w^24^. For each hemisphere, individual cortical thickness and T1/T2-weighted ratio maps from all HCP-YA participants—each represented as an M × N matrix, where M is the number of cortical vertices (samples) and N is the number of participants (features)—were concatenated along the column dimension, resulting in a combined matrix of shape M × 2N. Prior to conducting PCA, we applied Z-score normalization across vertices, thereby centering and scaling each feature to unit variance. PCA was then performed on this normalized matrix, treating cortical vertices as samples and the concatenated participant features as input variables. This decomposition produced a set of orthogonal principal components (PCs), consisting of spatial maps (i.e., PC scores) and corresponding participant-wise contributions (i.e., PC loading or individual weights) in pairs. The score map explains how the structural properties change across the cortical sheet on each component and the individual weights describe how individual cortical thickness and tissue density maps contribute to each component. The PCs are sorted in decreasing order according to the amount of variance explained by each of the components. The contribution ratio of density or thickness for a given PC was calculated by the ratio of the sum of absolute values of weights of the loading matrix from each measure to the sum of absolute values of weights from both measures.

#### The global hierarchy of architectural gradients

Two metrics which index the visual cortical functional hierarchy were used to validate the global hierarchy of the architectural gradients.

1. Population receptive field (pRF) size: It is widely known that pRF size progressively increases as one ascends the processing hierarchy from V1 to high-level visual cortex^33,46,47^. We first validated the global hierarchy of the architectural gradients by measuring if the gradients show similar spatial pattern to the pRF size across visual areas. Specifically, we calculated the Pearson correlation coefficients between architectural gradients and pRF size from the HCP’s 7T retinotopy dataset^45^. Only the vertices whose eccentricity of pRF within 8 degrees were used because retinotopic mapping stimuli were constrained to a circular region with a radius of 8 degrees.
2. Hierarchical rank: As the hierarchical level of the visual areas within the ventral stream has been widely studied and relatively clear, we reviewed literature describing the hierarchical relationships of 10 visual areas within the ventral stream^7,8,39,92–104^ and ranked them into a hierarchy from lowest-level to high: V1, V2, V3, V4, V8, PIT, VVC, FFC, TF, and PeEc, with cortical regions defined using labels of the HCP multimodal atlas^8^. Spearman’s rank correlation coefficients were then computed between the mean gradient score and the hierarchical rank of the areas.

#### Geometric models of the architectural gradients

The geometric models of the architectural gradients were constructed by the geodesic distance of each vertex to a set of specific references (Figure 1h). The primary gradient was modeled as the geodesic distance of each vertex of visual cortex to the calcarine sulcus (CS) anchor. The secondary gradient was modeled as the minimum geodesic distance from each vertex on the visual cortex to the eight anchors dispersed among four visual processing streams, which consisted of two local minima in the early visual cortex, two local minima in the dorsal visual stream, two local minima in the lateral visual stream, and two local minima in the ventral visual stream (Please see supplementary information for details, Table S2).

#### The gradient differences between vertices with different spatial distance

The gradient differences for each pair of vertices were calculated as the absolute difference between their gradient scores. The spatial distance between each pair of vertices was measured by the geodesic distance separating them on the cortical surface. The gradient differences between pairs of vertices were then sorted into 100 groups according to their spatial distance. The mean gradient differences and geodesic distances were then computed for each group and plotted against each other to evaluate how the gradient differences depend on spatial distance.

#### Functional significance of the architectural gradients

We characterized the functional significance of the architectural gradients by measuring to what extent each gradient is related to areal differentiation of the visual cortex^8^ and the fractional amplitude of low-frequency fluctuation (fALFF) from resting-state fMRI^51^.

1. Predicting visual areas based on the architectural gradients: Logistic Regression classifiers were trained on the architectural gradients to predict visual cortical areas. Specifically, the vertices from the visual cortex, with the architectural gradients as features, were used as the samples and the 44 visual cortical areas from the HCP MMP were used as the true class labels (i.e., 44-class classification). The areas with more vertices were down sampled to have the same number of vertices as the smallest areas to keep the number of samples with each class constant and thus avoid the imbalance of sample number across different visual areas. Logistic Regression classifiers were trained and tested on each downsampled data using a 5-fold cross-validation (CV) procedure. 100 random downsamples were performed and the averaged accuracy was used to measure the classification accuracy.
2. Predicting fALFF based on the architectural gradients: The fALFF is calculated as the ratio of the power spectrum of low-frequency modulations to that of the entire frequency range and is indicative of the magnitude of spontaneous brain activity^49,51^. For each rs-fMRI run, the time series of each vertex was Fourier transformed to a frequency domain without band-pass filtering, and the square root of the power was calculated at each frequency within the spectrum. fALFF was then computed by dividing the sum of amplitudes across a low-frequency band of the spectrum (0.01–0.1 Hz) by the sum of amplitudes across all frequencies up to the Nyquist frequency (0–0.625 Hz). The group fALFF map was calculated by averaging individual fALFF across all valid rfMRI runs within each participant and then across participants. 100 linear regression analyses using the 5-fold CV procedure were performed and the averaged R^2^ was used to characterize to what extent the two architectural gradients are related to spontaneous functional activity characterized by fALFF.

#### Laminar cytoarchitecture underlying the architectural gradients

The BigBrain cortical surfaces registered to standard surfaces, the surfaces of borders of the six cortical layers in BigBrain histological space, and the cell body density (CBD) data at 40-μm resolution were used together to extract laminar cytoarchitecture of the visual cortex. Specifically, for each of six cortical layers^90^, we first generated 10 surfaces based on the equivolumetric principle between its inner and outer borders^105^ to extract 10 CBD maps in the BigBrain space. Next, the CBD data from the BigBrain space were resampled to the fsLR_32k space, and then the 10 CBD maps were averaged to obtain an averaged CBD map for the layer. Finally, the spatial similarity between the architectural gradients and the averaged CBD map from each of the 6 cortical layers was measured with Pearson correlation coefficients.

#### Population receptive field (pRF) mapping

Functional images were preprocessed using a similar pipeline to the HCP data, including motion-correction, slice-timing correction, and phase-encoding distortion correction, and then aligned to each participant’s native cortical surface through FreeSurfer’s FS-FAST pipeline^106^. The preprocessed data from multiple runs from each participant were averaged to increase signal-to-noise ratio. The data were finally analyzed by a pRF model implemented in the VISTA Lab toolbox (github.com/vistalab), with additions for compressive nonlinearity (cvnlab.net/analyzePRF). The model predicts fMRI time series as the convolution of the stimulus-related time series and a canonical hemodynamic response function. The stimulus-related time series are in turn generated by computing the dot product between the stimulus apertures and a 2D isotropic Gaussian, scaling and applying a static power-law nonlinearity^107^. Several parameters of interest are produced from the pRF model for each vertex including phase angle, eccentricity, and pRF size. Vertices entered into any analyses presented in Figure 2 were only included if there was at least 10% variance-explained by the model-fit.

To produce maps of visual field coverage (Figure 2h) similar to previous work^34^, each participant’s pRF fits from the anterior temporal (AT) region were simulated as 2-D gaussians and plotted on the visual field with a maximum value of 1, summing across pRF’s within a participant without allowing values at any given point in the visual field to exceed 1. PRF fits with sigma values smaller than 0.5 or eccentricity values larger than the stimulus-mapping field-of-view (11.5 degrees of visual angle) were excluded. The coverage map for a given participant’s AT region was bootstrapped 1000 times, plotting 75% of pRF fits on each iteration, then averaged across bootstraps. Coverage maps were then averaged across participants to produce Figure 2h.

To quantify whether there is a left-right contralateral bias in the sampling of visual space (and to test whether such a bias is significantly different in each hemisphere), we calculated for each pRF a laterality index as previously defined by Sheremata and Silver (2015)^58^ according to the equation below:

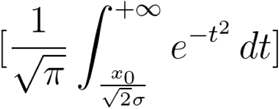

Where resulting values of 1 mean the pRF is contralateral, 0.5 is no laterality bias, and 0 is ipsilateral bias. Additionally, we input pRF sigma values that were adjusted for the non-linearity exponent as defined by Kay et al. (2013)^107^. For the purposes of visual comparison, we subtracted 0.5 from index values so that resulting laterality scores were relative to 0 to represent the center of the visual field, and then values were inverted with a -1 scalar so that left hemisphere pRF laterality index values are plotted on the right side of space, and the right hemisphere on the left as shown in Figure 2i. The laterality index was calculated for each pRF for a given participant and then averaged within that participant to result in a single mean laterality index for the left hemisphere pRFs and a single index for their right hemisphere pRFs. The histograms illustrated in Figure 2i depict density of participants (kernel smoothed).

#### Behavioral relevance of the architectural gradients

The behavioral significance of the architectural gradients was examined by measuring how the individual weights from PCA can account for the individual variation in behavioral performance on related visual tasks. The HCP-YA dataset include behavioral tasks of a range of motor, sensory, cognitive, and emotional processes. Because our architectural gradients cover the entire visual cortex and may involve into a variety of cognitive and behavioral abilities, a total of 15 vision-related or vision-based behavioral tasks^60^, designed to measure nonverbal episodic memory ability, cognitive flexibility, inhibitory control and attention ability, fluid intelligence, reading decoding skill, general vocabulary knowledge, speed of processing, spatial orientation processing ability, sensitivity of sustained attention, specificity of sustained attention, verbal episodic memory ability, working memory ability, odor identification ability, visual acuity, and contrast sensitivity, were selected to examine the behavioral relevance of the gradients (Please see supplementary information for details, Table S3).

For the two sets of individual weights (i.e., myelin/neurite density weights and cortical thickness weights) associated with each of architectural gradient, we carried out a single integrated multivariate analysis using canonical correlation analysis (CCA)^59^, to simultaneously co-analyze the two sets of weights along with 15 behavioral variables from all participants. CCA aims to identify symmetric linear relations between the two sets of variables. That is, we used CCA to find components that relate the two sets of weights from each gradient to 15 sets of participants’ behavioral measures. Each CCA component identifies a linear combination of two weights and a linear combination of behavioral variables, where the variation in strength of involvement across participants is maximally correlated. The normalized magnitude of behavioral weights, obtained by dividing each squared behavioral weight by the total sum of all squared weights, was used to characterize the behavioral profile of an architectural gradient and reveal how the two gradients are different in predicting the same set of behavioral variables.

#### The development of the architectural gradients across the lifespan

We divided HCP-D, HCP-YA, and HCP-A participants into different age windows (subgroups) in ascending order of age. Among them, the participants of HCP-D and HCP-A were sorted by their age in months, while HCP-YA sorted in years because no month information was provided. Each window consisted of 50 participants and had a step size of ten. As a result, there are 61, 103, and 68 age windows generated for HCP-D, HCP-YA, and HCP-A data, respectively. To characterize the changes of the architectural gradients across the human lifespan, we conducted PCA on the stacked tissue density and thickness maps of each window as we did in the whole HCP-YA data. We label the principal components from each age window as lifespan components (LC) to differentiate them from the original PCs derived from the HCP-YA dataset. To determine if the observed LCs were similar to the PCs from the HCP-YA data, we calculated the Pearson’s correlation coefficients between the HCP-YA PCs and the top ten LCs for each window. The LC which shows strongest correlation with a PC was considered to be the correspondence LC to the PC in that age window. Lastly, how the observed gradients from each age window change relative to the gradient from the HCP-YA was measured by the Pearson’s correlation coefficient between their score maps. The changes along age were then charted (i.e., developmental trajectories). A linear model was then constructed to characterize the developmental trajectory within each of three datasets. The slope of the linear model was defined as the annualized rate of change (AROC) in the architectural gradients.

## Ethical Compliance

Data analyzed from the Human Connectome Project follows all necessary privacy and security guidelines. Data collected from participants at Princeton University followed all Institutional Review Board ethics and guidelines (protocol number 13074), and all safety regulations set forth by the Scully Center for the Neuroscience of Mind and Behavior. Informed consent was collected from every participant involved in this study, and all were reimbursed for their time.

## Data Availability

The data from the HCP Young Adult (HCP-YA) are publicly available at the https://www.humanconnectome.org; The data from the HCP Development (HCP-D) and HCP Aging (HCP-A) are publicly available at https://nda.nih.gov. Users can access these after registration.

## Code Availability

The HCP data were preprocessed using the HCP-Pipeline analyses (https://github.com/Washington-University/HCPpipelines). Custom code for gradient analysis can be found at https://github.com/BNUCNL/VisualCortexGradient. Code to reproduce population receptive field mapping figures can be found at: https://github.com/gomezj/entorhinal_prf.

## Acknowledgements

We thank Youyi Liu and the members of the Brain Development Lab for their constructive discussions. This work was supported by Brain Science and Brain-like Intelligence Technology - National Science and Technology Major Project (2021ZD0200500) to ZZ and by start-up funds from the Princeton Neuroscience Institute to JG.

## Author Contributions

JG and ZZ conceived of the idea and design of the study. XC and XL compiled the data, performed the analyses, and prepared visualizations. PH, ED, and JY performed the analyses. JG, XC, XL, and ZZ drafted and revised the manuscript with input from all authors. JG and ZZ supervised the research.

