## Supplemental Texts, Figures and Tables for "Distinct gradients of cortical architecture capture visual representations and behavior across the lifespan"

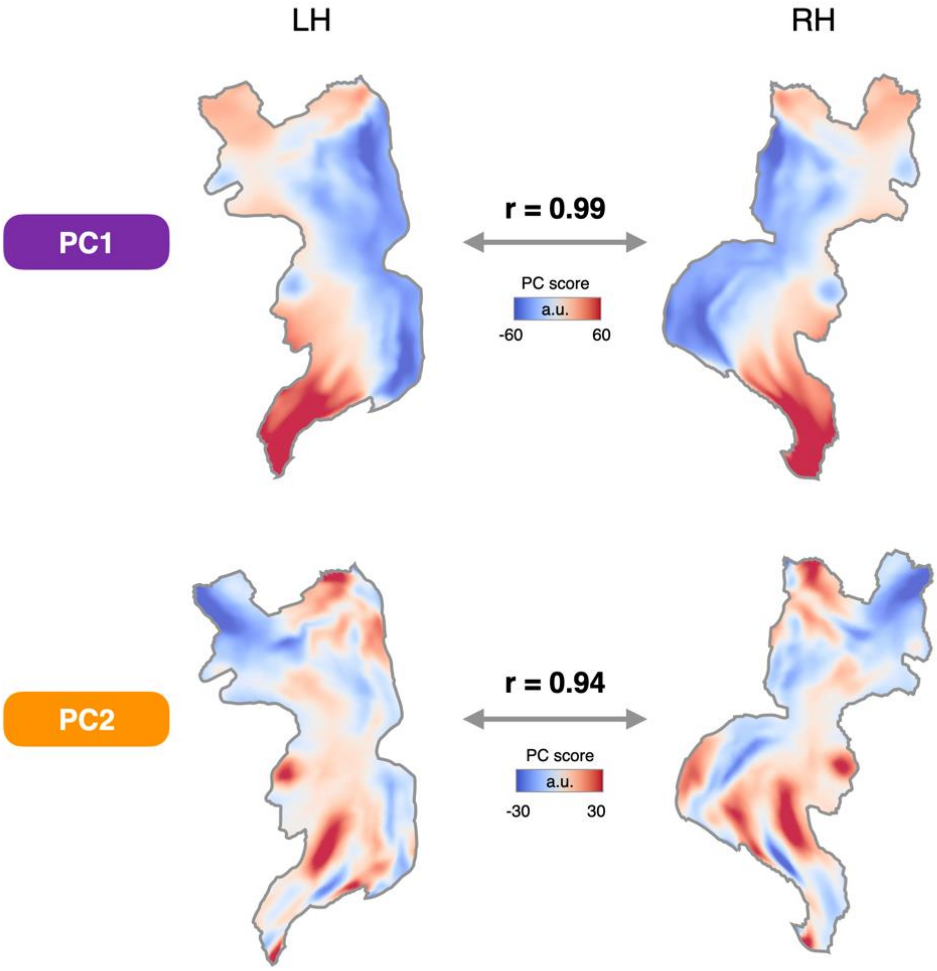


**Figure S1: The two hemispheres have very similar architectural gradients in visual cortex; related to Figure 1.** The two architectural gradients from the principal component analysis (PCA) on the left hemisphere (left penal) and the right hemisphere (right panel). The gradients show high spatial similarity between two hemispheres as measured with a vertex-wise Pearson correlation (gradient 1: r = 0.99; gradient 2: r = 0.94).

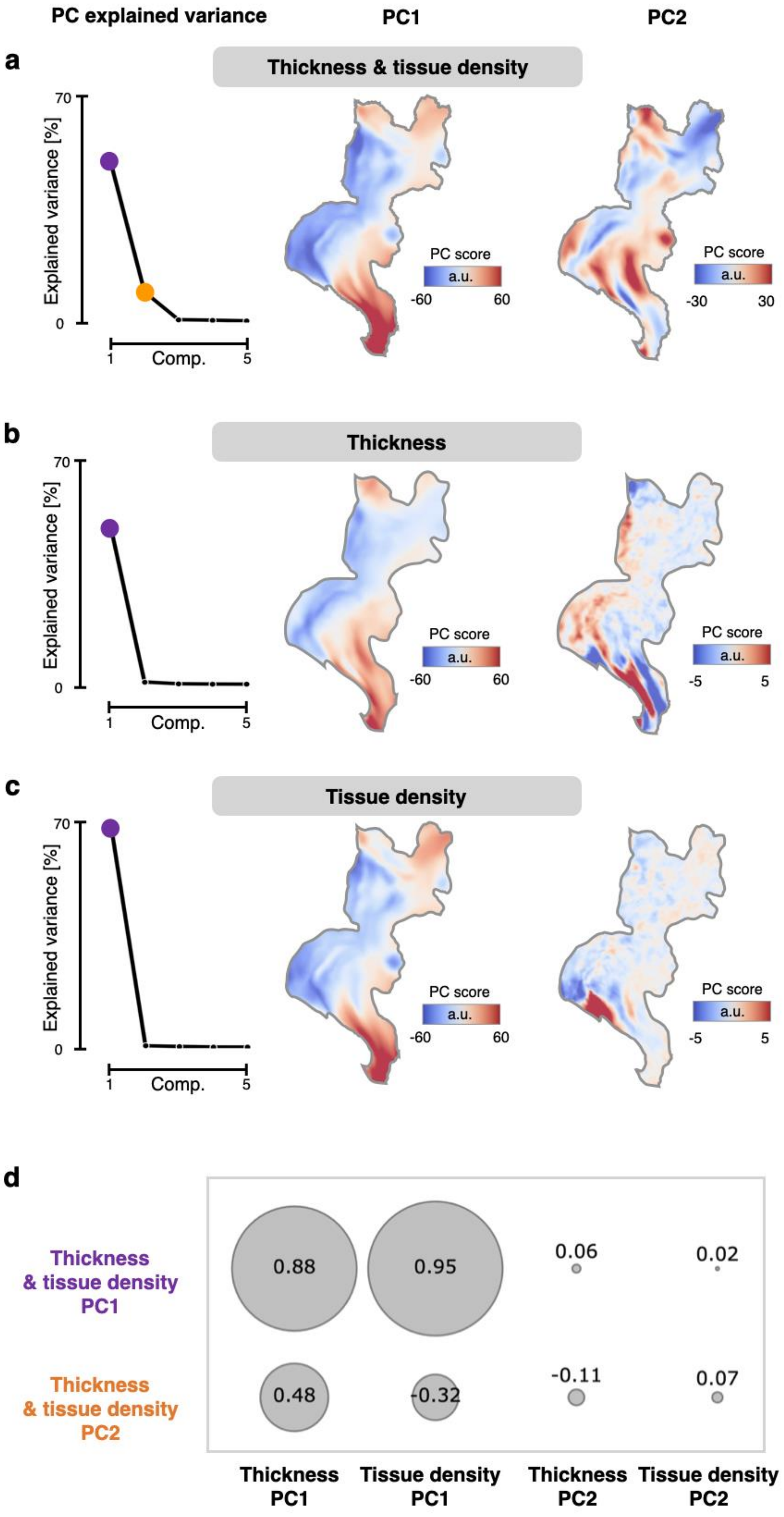
PC explained variance
PC1
PC2
a
Thickness & tissue density
70
Explained variance [%]
0
1
Comp.
5
PC score
a.u.
-60
60
PC score
a.u.
-30
30
b
Thickness
70
Explained variance [%]
0
1
Comp.
5
PC score
a.u.
-60
60
PC score
a.u.
-5
5
c
Tissue density
70
Explained variance [%]
0
1
Comp.
5
PC score
a.u.
-60
60
PC score
a.u.
-5
5
d
Thickness & tissue density PC1
0.88
0.95
0.06
0.02
Thickness & tissue density PC2
0.48
-0.32
-0.11
0.07
Thickness PC1
Tissue density PC1
Thickness PC2
Tissue density PC2

**Figure S2: The principal component analysis (PCA) on the concatenated cortical thickness and tissue density maps can exploit the concurrent spatial changes of the two measurements and thus extract more meaningful architectural gradients than PCA on a single architectural measurement; related to Figure 1. (a)** The gradients from the PCA performed on the concatenated cortical thickness and tissue density maps. **(b)** The gradients from the PCA using only cortical thickness maps. **(c)** The gradients from the PCA using only tissue density (T1w/T2w ratio) maps. **(d)** The spatial pattern similarities between the gradients from the concatenated PCA and the separate PCA on the single architectural measurement, measured by Pearson correlation coefficients. While the separate PCAs on the single measurements can extract similar PC1s to the concatenated PCA, they fail to extract a meaningful PC2.

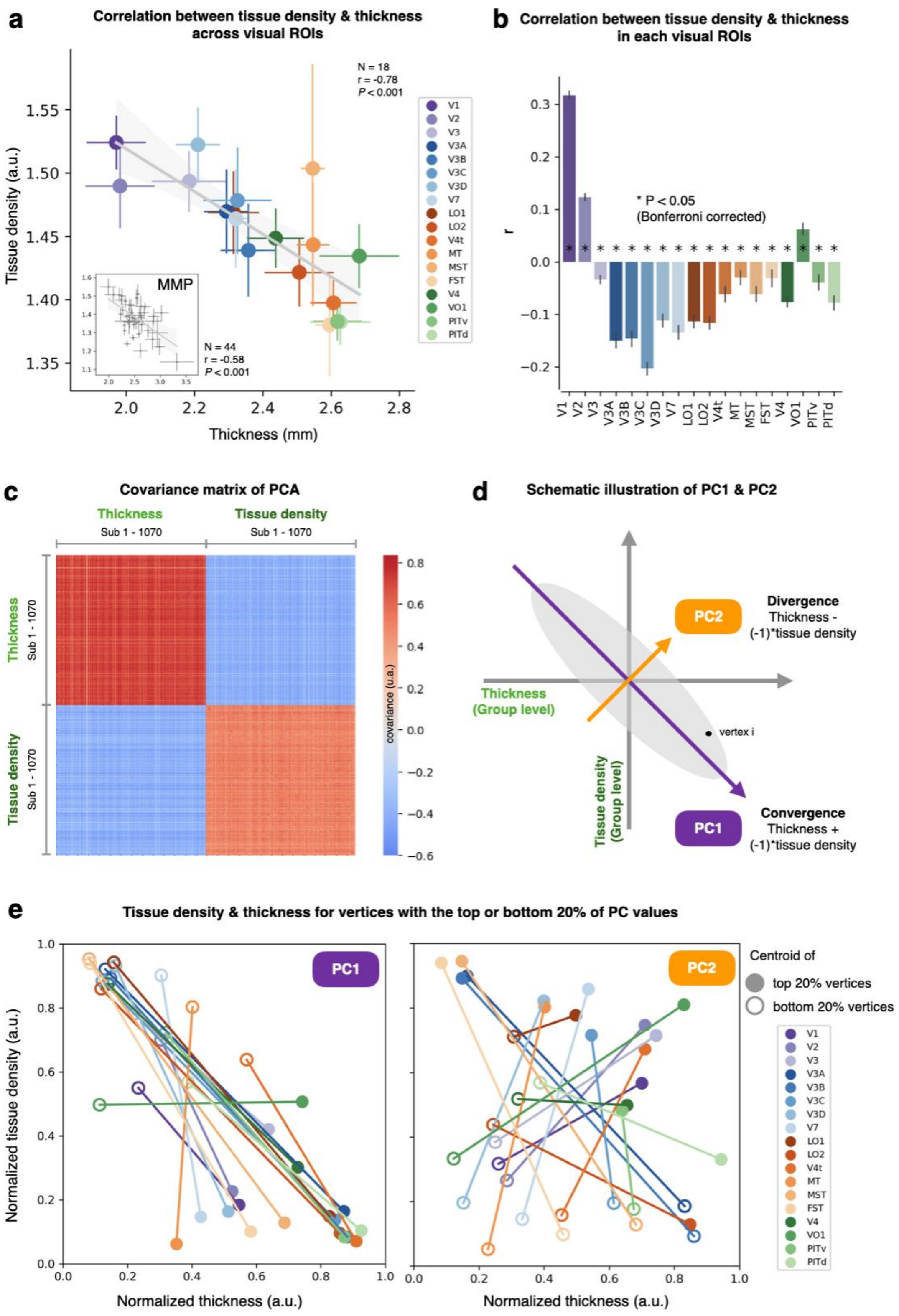


**Figure S3: PCA and the convergent and divergent patterns between group average maps of cortical thickness and tissue density; related to Figure 1. (a)** Relationship between cortical thickness and tissue density across large-scale visual ROIs. Parcellations were defined using an independent dataset (Abdollahi et al., 2014). Each colored point represents the mean values within a visual area, with error bars indicating the SD. The inset shows the same analysis using visual ROIs from the MMP atlas (Glasser et al., 2016). Strong negative correlations were observed across visual ROIs in both parcellations. **(b)** Relationship between cortical thickness and tissue density within individual visual ROIs. Bar plots show correlation coefficients (r) computed separately for each ROI, with error bars indicating the SEM across hemispheres. Most visual areas showed significant negative correlations (*$P < 0.05$, Bonferroni corrected), whereas V1, V2, and VO1 displayed positive correlations, consistent with previous findings (Maingault et al., 2021; Sereno et al., 2013; Shafee et al., 2015). **(c)** The covariance matrix of the PCA reveals high inter-individual positive covariance within cortical thickness and tissue density, respectively, and inter-individual negative covariance between cortical thickness and tissue density. This suggests that the group-level average thickness and tissue density account for a significant portion of the variance, allowing the high-dimensional features to be approximated as two primary dimensions: group-level thickness and tissue density. **(d)** Schematic illustration of PC1 and PC2. As indicated by (c), the PCA can be approximated as a two-dimensional problem. The point cloud, represented by the gray ellipse, indicates individual vertices. The principal component with the greatest explained variance (PC1) is oriented along thickness + (-1) * tissue density, while the PC2 is oriented along thickness - (-1) * tissue density. **(e)** Alignment of PC scores with local microstructural features. Within each visual ROI, mean cortical thickness and density were plotted for vertices with the top and bottom 20% of PC scores (closed and open dots, respectively), connected by lines. For PC1 (left), vertices with higher scores generally showed greater thickness and lower density, while those with lower scores showed the opposite trend. This within-ROI relationship parallels the large-scale cross-ROI anti-correlation in (a), consistent with PC1 reflecting a large-scale cortical organizational axis. In contrast, PC2 (right) exhibited more heterogeneous profiles across ROIs, indicating that it captures region-specific microstructural variations.

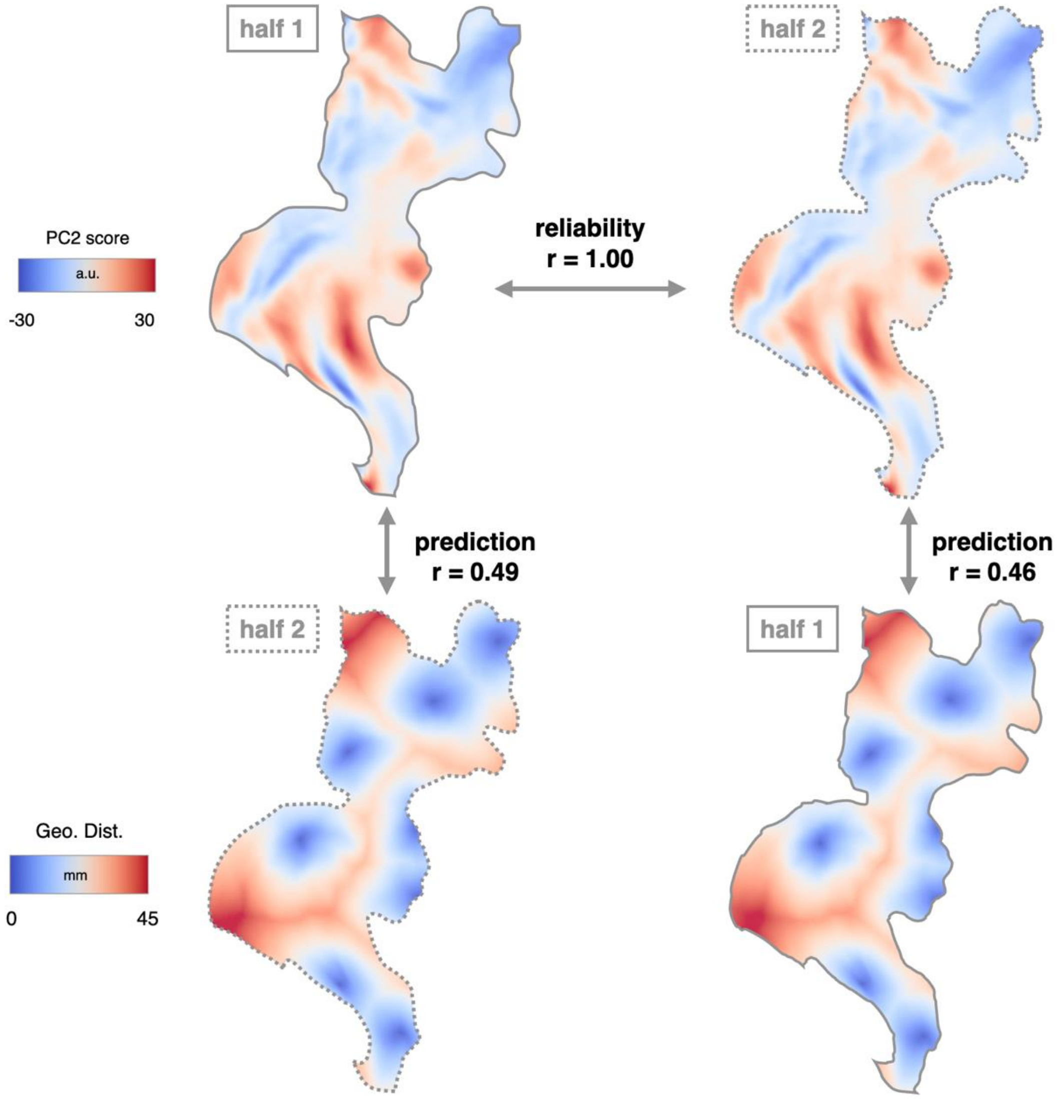


**Figure S4: Split-half cross-validation of PC2 maps demonstrates reliable geometric features underlying visual cortex organization; related to Figure 1.** A split-half cross-validation was conducted to mitigate concerns about circularity in defining the geodesic model's anchor points. Anchors were defined from one half of the participants and used to predict the PC2 map in the other half. Top row shows the PC2 maps derived from half 1 (left, solid outline) and half 2 (right, dashed outline) of the sample, displaying highly similar spatial patterns ($r = 1.00$, $p < 0.001$), indicating strong reliability. Middle and bottom rows demonstrate cross-prediction: the geodesic model derived on half 2 successfully predicts the PC2 map in half 1 (left, $r = 0.49$), and vice versa (right, $r = 0.46$). These results indicate reliable geometric features underlying the spatial organization of PC2 across independent samples.

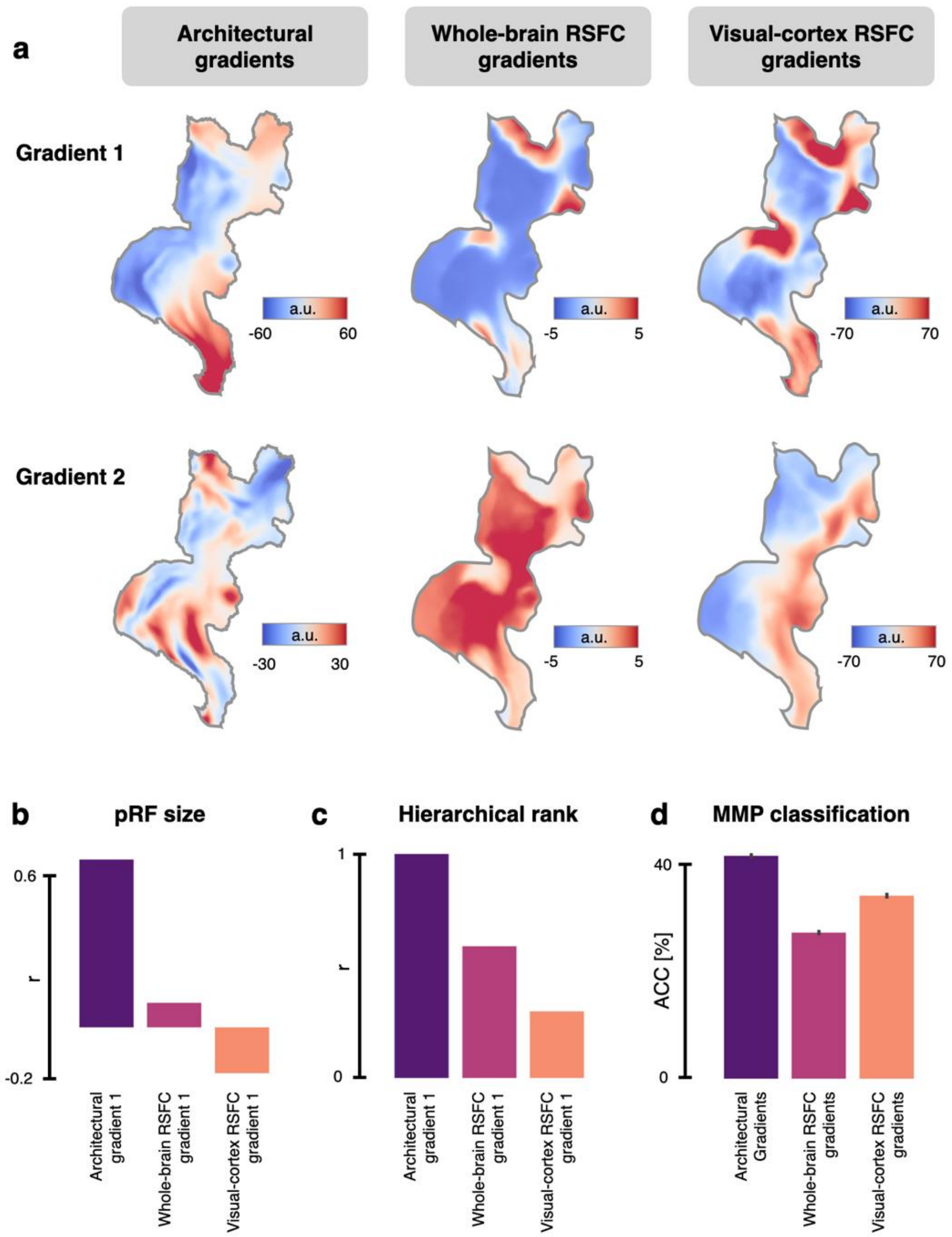


**Figure S5: The architectural gradients derived from the principal component analysis (PCA) on the mesoscale structural measurements are more functionally relevant than the gradients from resting-state functional connectivity (RSFC); related to Figure 2. (a)** The first two gradients from the PCA on the mesoscale structural measurements (left), the results from diffusion embedding on the whole-brain RSFC matrix (middle) (Margulies et al., 2016), and the PCA on the RSFC matrix limited to include only visual cortex (right). **(b-c)** Architectural gradient 1 is more related to visual processing hierarchy than either of the primary gradient from the whole-brain RSFC or visual-cortex RSFC, which was evaluated based on a map's correlation to pRF size (panel b) and hierarchical rank (panel c). **(d)** The first two architectural gradients are more powerful in classifying visual areas than the two gradients from the whole-brain RSFC and visual-cortex RSFC.

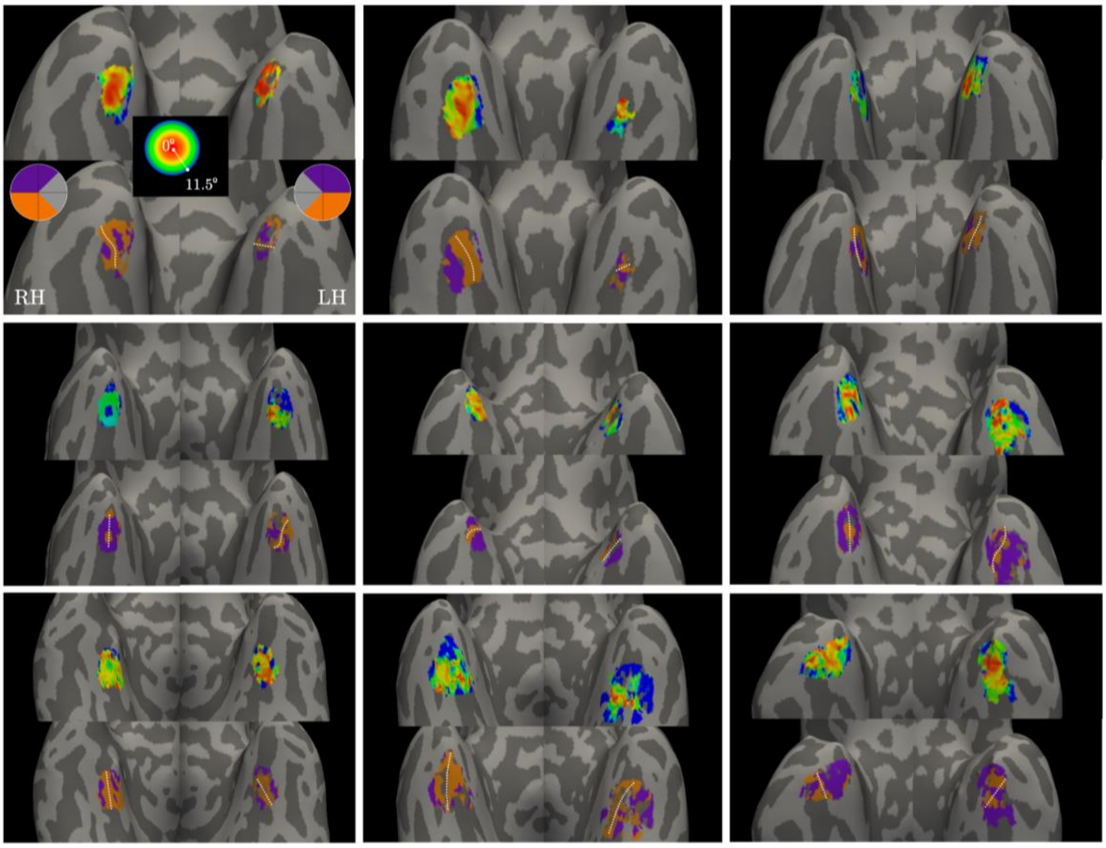


**Figure S6: Anterior temporal visual representation clusters in all participants; related to Figure 2.** Individual cortical surfaces have been inflated and zoomed to make visible the ventral surface of the anterior temporal pole. Vertices are colored by properties of the fitted pRF model, showing either eccentricity (top panel in each participant) or polar angle (bottom panels). Color scales denoting eccentricity and polar angle value mappings shown in the inset of P1. White circles outline the putative AT visual representation cluster, and dotted white lines are drawn through the shared lower visual field representation in polar angle maps, bisecting the pRF cluster into two hemifield representations. Visualized data is thresholded, showing only vertices whose time course was fitted by the pRF model with at least 10% variance explained.

| Area Name | Area Description | Other Names | Key Studies |
|---|---|---|---|
| V1 | Primary Visual Cortex | 17, hOC1, OC, BA17 | Amunts et al 2000, Fischl et al 2008, Abdollahi et al 2014 |
| MST | Medial Superior Temporal Area | MSTv, hOC5, hOC5v | Abdollahi et al 2014, Kolster et al 2010, Malikovic et al 2007, Fischl et al 2008 |
| V6 | Sixth Visual Area | 112 | Pitzalis et al 2006, Pitzalis et al 2013, Sereno et al 2012, Nieuwenhuys et al 2014 |
| V2 | Second Visual Area | 18, hOC2, OB, BA18 | Amunts et al 2000, Fischl et al 2008, Schira et al 2009, Abdollahi et al 2014, Wang et al 2015, Wandell and Winawer 2011 |
| V3 | Third Visual Area | V3d, V3v, VP, hOC3d, hOC3v | Abdollahi et al 2014, Rottschy et al 2007, Schira et al 2009, Kujovic et al 2012, Wang et al 2015, Wandell and Winawer 2011 |
| V4 | Fourth Visual Area | V4d, V4v, hV4, hOC4v, hOC4lp, LO1 | Hansen et al 2007, Abdollahi et al 2014, Rottschy et al 2007, Malikovic et al 2015 |
| V8 | Eighth Visual Area | VO1 | Hadjikhani et al 1998, Abdollahi et al 2014 |
| V3A | Area V3A | V3D, hOC4d | Abdollahi et al 2014, Swisher et al 2007, Kujovic et al 2012, Wandell and Winawer 2011, Larsson and Heeger 2006, Tootell et al 1997 |
| POS2 | Parieto-Occipital Sulcus Area 2 | | Glasser and Van Essen 2011 |
| V7 | Seventh Visual Area | IPS0 | Abdollahi et al 2014, Swisher et al 2007, Larsson and Heeger 2006, Tootell et al 1998, Hagler et al 2007, Wang et al., 2015 |
| IPS1 | IntraParietal Sulcus Area 1 | | Swisher et al 2007, Wang et al., 2015, Hagler et al 2007 |

| | | | |
|---|---|---|---|
| FFC | Fusiform Face Complex | FFA, FG2 | Glasser and Van Essen 2011, Kanwisher and Yovel, 2006, Caspers et al 2013, Weiner et al 2014 |
| V3B | Area V3B | V3C | Abdollahi et al 2014, Larsson and Heeger 2006, Swisher et al 2007, Wandell and Winawer 2011, Smith et al 1998 |
| LO1 | Area Lateral Occipital 1 | LO2, hOC4la | Abdollahi et al 2014, Hansen et al 2007, Malikovic et al 2015, Larsson and Heeger 2006 |
| LO2 | Area Lateral Occipital 2 | LO1, hOC4la | Abdollahi et al 2014, Hansen et al 2007, Malikovic et al 2015, Larsson and Heeger 2006 |
| PIT | Posterior InferoTemporal Complex | phPITv, phPITd, OFA, hOC4la | Abdollahi et al 2014, Kolster et al 2010, Malikovic et al 2015, Kanwisher and Yovel, 2006, Tsao et al 2008 |
| MT | Middle Temporal Area | hOC5, hOC5d | Abdollahi et al 2014, Kolster et al 2010, Malikovic et al 2007, Fischl et al 2008 |
| PCV | PreCuneus Visual Area | PrCu | Sereno et al 2012 |
| 7Pm | Medial Area 7P | 7P | Scheperjans et al 2008a, Scheperjans et al 2008b |
| POS1 | Parieto-Occipital Sulcus Area 1 | "Retrosplenial Cortex" | Glasser and Van Essen 2011 |
| 7Am | Medial Area 7A | | Scheperjans et al 2008a, Scheperjans et al 2008b |
| 7PL | Lateral Area 7P | | Scheperjans et al 2008a, Scheperjans et al 2008b |
| LIPv | Area Lateral IntraParietal ventral | hIP3 | Van Essen et al 2012a, Scheperjans et al 2008a, Scheperjans et al 2008b |
| VIP | Ventral IntraParietal Complex | | Van Essen et al 2012a |
| MIP | Medial IntraParietal Area | | Van Essen et al 2012a |
| ProS | ProStriate Area | | Glasser and Van Essen 2011, Vogt et al 2001, Sanides and Vitzthum 1965, Sanides, 1970 |

| | | | |
|---|---|---|---|
| PeEc | Perirhinal Ectorhinal Cortex | ATFP, AFP1, Ectorhinal, Perirhinal, 35, 36 | Augustinack et al 2013, Ding et al 2009, Ding and Van Hoesen 2010, Rajimehr et al 2009, Tsao et al 2008 |
| PHA1 | ParaHippocampal Area 1 | | |
| PHA3 | ParaHippocampal Area 3 | | |
| TF | Area TF | | von Economo and Koskinas 1925, Triarhou 2007 |
| PH | Area PH | | von Economo and Koskinas 1925, Triarhou 2007 |
| DVT | Dorsal Transitional Visual Area | | |
| IP1 | Area IntraParietal 1 | | Choi et al 2006 |
| IP0 | Area IntraParietal 0 | | |
| V6A | Area V6A | 112 | Pizalis et al 2013, Nieuwenhuys et al 2014 |
| VMV1 | VentroMedial Visual Area 1 | PHC2, PHC-2 | Arcaro et al 2009, Wang et al 2015 |
| VMV3 | VentroMedial Visual Area 3 | VO2 | Arcaro et al 2009, Wang et al 2015, Wandell and Winawer 2011 |
| PHA2 | ParaHippocampal Area 2 | | |
| V4t | Area V4t | LO2 | Abdollahi et al 2014, Kolster et al 2010, Larsson and Heeger 2006 |
| FST | Area FST | | Abdollahi et al 2014, Kolster et al 2010 |
| V3CD | Area V3CD | V3A, V3B, hOC4la | Abdollahi et al 2014, Malikovic et al 2015 |
| LO3 | Area Lateral Occipital 3 | hOC4la | |
| VMV2 | VentroMedial Visual Area 2 | PHC1, PHC-1 | Arcaro et al 2009, Wang et al 2015 |
| VVC | Ventral Visual Complex | VO1, VO2, PHC1, PHC2, PHC-1, PHC-2, FG1 | Arcaro et al 2009, Wang et al 2015, Wandell and Winawer 2011, Caspers et al 2013, Weiner et al 2014 |

**Table S1: Detailed descriptions of the 44 visual areas from the HCP multimodal parcellation (MMP) atlas; related to Figure 1.**

| Hemisphere | Visual Stream | x | y | z | Anatomical Landmark |
|---|---|---|---|---|---|
| Left | Early | -7.22876 | -96.1693 | 10.94631 | Upper bank of the Calcarine Sulcus |
| | Early | -10.1512 | - | -11.9219 | Lower bank of |

| | | | | | |
|---|---|---|---|---|---|
| | | | 92.6749 | | the Calcarine Sulcus |
| | Dorsal | -11.0877 | -55.9231 | 62.04463 | Intersection between Superior Parietal Gyrus and Precuneus. |
| | Dorsal | -14.3971 | -75.7594 | 43.54079 | Confluence of Superior Occipital Gyrus, Parieto-Occipital Sulcus and Superior Parietal Gyrus |
| | Lateral | -43.357 | -69.2016 | -2.55437 | Intersection between Anterior Occipital Sulcus and Inferior Occipital Gyrus |
| | Lateral | -41.69 | -76.4999 | 16.17956 | Intersection between Middle Occipital Sulcus and Middle Occipital Gyrus |
| | Ventral | -30.1495 | -32.6623 | -16.5016 | Within Collateral Sulcus |
| | Ventral | -40.1003 | -19.7558 | -23.9135 | Within anterior transverse Collateral Sulcus |
| Right | Early | 11.34271 | -94.6688 | 13.01122 | Upper bank of the Calcarine Sulcus |
| | Early | 8.911536 | -87.6605 | -10.5582 | Lower bank of the Calcarine Sulcus |
| | Dorsal | 11.16673 | -55.0289 | 62.86484 | Intersection between Superior Parietal Gyrus and Precuneus. |
| | Dorsal | 17.84735 | -73.4929 | 44.53817 | Confluence of Superior Occipital Gyrus, Parieto-Occipital Sulcus and Superior Parietal Gyrus |
| | Lateral | 45.97639 | -66.362 | -5.06974 | Intersection between Anterior Occipital Sulcus and Inferior Occipital Gyrus |

| | | | | | |
|---|---|---|---|---|---|
| | Lateral | 44.79291 | -<br>72.1836 | 15.25219 | Intersection between Middle Occipital Sulcus and Middle Occipital Gyrus |
| | Ventral | 31.54866 | -<br>29.0792 | -18.4202 | Within Collateral Sulcus |
| | Ventral | 38.7923 | -<br>9.02916 | -32.9193 | Within anterior transverse Collateral Sulcus |

**Table S2: Location information of anchors of the secondary gradient's geometric model; related to Figure 1.**

| Full Display Name | Variable | Assessment | Description |
|---|---|---|---|
| NIH Toolbox Picture Sequence Memory Test: Unadjusted Scale Score | PicSeq_Unadj | Nonverbal Episodic Memory | The picture sequence is presented on a computer screen in a particular order, while the content of the pictures is described through audio. After presentation, the sequence of pictures is scrambled and the participant is asked to move the pictures to restore the correct order (Dikmen et al., 2014). |
| NIH Toolbox Dimensional Change Card Sort Test: Unadjusted Scale Score | CardSort_Unadj | Cognitive Flexibility | The participant is required to select a test picture (e.g. white rabbit and brown sailboat) that match the target picture (e.g. brown rabbit) based on a specified dimension (e.g. color or shape). |
| NIH Toolbox Flanker Inhibitory Control and Attention Test: Unadjusted Scale Score | Flanker_Unadj | Inhibitory Control and Attention | The participant is required to focus on a given stimulus while inhibiting attention to stimuli flanking it. Sometimes the middle stimulus is pointing in the same direction as the flankers (congruent) and sometimes in the opposite direction (incongruent). |
| Penn Progressive Matrices: Number of Correct Responses | PMAT24_A_CR | Fluid Intelligence | Based on an abbreviated version of the Raven's Progressive Matrices Form A developed by Gur and colleagues (Bilker et al. 2012). The participant is presented with an array of pictures arranged in a 2x2, 3x3 or 1x5 grid, with one cell left empty. The participant is required to reason based on the pictures presented and select the most suitable option to fill in the empty cell from five choices. |
| NIH Toolbox Oral Reading Recognition Test: Unadjusted Scale Score | ReadEng_Unadj | Reading Decoding | A single word or letter is presented on a computer screen, and the participant is asked to read it aloud. An examiner then judges whether the pronunciation is correct (Gershon et al., 2013). |

| | | | |
|---|---|---|---|
| NIH Toolbox Picture Vocabulary Test: Unadjusted Scale Score | PicVocab_Unadj | Vocabulary Comprehension | The participant is asked to listen to a word, and then selects the picture that best corresponds to the meaning of the word from four pictures presented on the screen. The pictures depict objects, actions, and conceptual descriptions (e.g. ball, running, friendship) (Gershon et al., 2013). |
| NIH Toolbox Pattern Comparison Processing Speed Test: Unadjusted Scale Score | ProcSpeed_Unadj | Processing Speed | Two pictures are presented simultaneously and the participant is asked to judge whether they are the same or not. The task uses simple pictures to measure processing speed as purely as possible. |
| Variable Short Penn Line Orientation: Total Number Correct | VSPLOT_TC | Spatial Orientation Processing | Two line segments (blue and red) are presented on the screen, and the participant is asked to rotate the blue segment using keyboard buttons to make it parallel with the red segment. |
| Short Penn Continuous Performance Test: Sensitivity = SCPT_TP/(SCPT_TP + SCPT_FN) (SCPT_SEN) | SCPT_SEN | Sustained Attention Sensitivity | The participant is asked to look at the flashing vertical and horizontal red lines on the computer screen. In one condition, the participant must press the space bar when the lines form a number, and in another condition, the participant must press the space bar when the lines form a letter. |
| Short Penn Continuous Performance Test: Specificity = SCPT_TN/(SCPT_TN + SCPT_FP) (SCPT_SPEC) | SCPT_SPEC | Sustained Attention Specificity | |
| Penn Word Memory Test: Total Number of Correct Responses | IWRD_TOT | Verbal Episodic Memory | This task presents 20 words for participants to memorize. In the subsequent memory test, 40 words are presented, including the previous 20 words and 20 new words. Participants are required to judge whether they have seen the words before, with four options to choose from: |

| | | | |
|---|---|---|---|
| | | | "definitely yes", "probably yes", "probably no", and "definitely no". |
| NIH Toolbox List Sorting Working Memory Test: Unadjusted Scale Score | ListSort_Unadj | Working Memory | This task requires the participant to sequence visually- and orally-presented stimuli (foods and animals) into size order. |
| NIH Toolbox Odor Identification Age 3+ Unadjusted Scale Score | Odor_Unadj | Odor Identification | This task requires the participant to identify which of the four presented pictures matches the odor they just smelled. |
| EVA score - Denominator | EVA_Denom | Visual Acuity | Based on the Snellen chart, the participant with corrected vision is asked to read letters that gradually become smaller. |
| Mars Final Contrast Sensitivity Score | Mars_Final | Contrast Sensitivity | This task requires the participant to read letters on the Mars Contrast Test Chart from left to right and top to bottom, as the contrast of the letters gradually decreases, until making two consecutive errors and stopping the test. |

**Table S3: Detail information of 15 vision-related or vision-based behavioral tasks selected from HCP-YA dataset; related to Figure 3.**
